# mobileOG-db: a manually curated database of protein families mediating the life cycle of bacterial mobile genetic elements

**DOI:** 10.1101/2021.08.27.457951

**Authors:** Connor L. Brown, James Mullet, Fadi Hindi, James E. Stoll, Suraj Gupta, Minyoung Choi, Ishi Keenum, Peter Vikesland, Amy Pruden, Liqing Zhang

**Affiliations:** Genetics, Bioinformatics, and Computational Biology, Virginia Tech, Blacksburg, VA, 24061, USA; Civil and Environmental Engineering, Virginia Tech, Blacksburg, VA, 24061, USA; Fralin Life Science Institute, Blacksburg, VA, 24061, USA; Computer Science, Virginia Tech, Blacksburg, VA, 24061, USA

**Author notes:** To whom correspondence should be addressed. Tel: 1-540/231-6635 Fax: 540/231-7532 Fax: 540/231-7532.

## Abstract

Currently available databases of bacterial mobile genetic elements (MGEs) contain both “core” and accessory MGE functional modules, the latter of which are often only transiently associated with the element. The presence of these accessory genes, which are often close homologs to primarily immobile genes, limits the usability of these databases for MGE annotation. To overcome this limitation, we analysed 10,776,212 protein sequences derived from seven MGE databases to compile a comprehensive database of 6,140 manually curated protein families that are linked to the “life cycle” (integration, excision, replication/recombination/repair, transfer, and stability/defense) of all major classes of bacterial MGEs. We overlay experimental information where available to create a tiered annotation scheme of high-quality annotations and annotations inferred exclusively through bioinformatic evidence. We additionally provide an MGE-class label for each entry (e.g., plasmid, integrative element) derived from the source database, and assign a list of keywords to each entry to delineate different MGE functional modules and to facilitate annotation. The resulting database, mobileOG-db (for mobile orthologous groups), provides a simple and readily interpretable foundation for an array of MGE-centred analyses. mobileOG-db can be accessed at mobileogdb.flsi.cloud.vt.edu/, where users can browse and design, refine, and analyse custom subsets of the dynamic mobilome.

## INTRODUCTION

Prokaryotic genomes undergo frequent interactions with mobile genetic elements (MGEs) including plasmids, bacteriophages, insertion sequences, and other integrative elements (IGEs). These interactions can confer beneficial or detrimental properties to the organism (1), and in some cases appear to have little impact on the organism at all (2). Of particular importance to public health, bacterial MGEs can function as key vehicles for the proliferation of antimicrobial resistance (AMR), which is now pandemic in many clinically-significant bacteria (3). Emerging efforts aimed at developing frameworks for predicting the emergence of novel antibiotic resistance genes (ARGs), whose products confer AMR in bacteria, have identified associations with MGEs as a key indicator of a potential novel ARG (4).

While there are many tools and databases available for annotating mobile genetic elements (7–10), there is at present no centralized resource for mobile genetic element “hallmark” genes which can serve as the basis for annotating diverse classes of MGEs, such as is aggregated by pVOG (11) for phages. However, even pVOG includes many primarily cellular genes (11, 12), which they identify based on the frequency of occurrence in phages relative to the occurrence in cellular genome sequences. Similarly, public databases of full length mobile genetic elements, such as ACLAME (6) or ICEBerg (13), comprise both core and accessory MGE genes. While such databases are highly informative, the presence of these cargo genes leads to frequent occurrences of false-positive hits that confound and complicate the annotation of MGEs (14). In sum, the presence of these cargo genes creates the need for extensive expertise and research to detect, analyse, and annotate diverse types of mobile genetic elements in biological data. his is a key obstacle particularly for antibiotic resistance research, where knowledge of the carriage of ARGs on MGEs is highly valuable towards identifying mobile ARGs. (15).

To facilitate MGE annotation, we propose the mobile orthologous groups database and webserver (mobileOG-db), an interactive resource compiling knowledge encompassing a comprehensive variety of proteins mediating the essential functions of all major classes of bacterial MGEs. Here we define the essential functions or “life cycle” of MGEs as (1) integration and excision (IE) from one genetic locus to another; (2) replication, recombination, or repair (RRR) of element nucleic acid; (3) inter-organism transfer (T); (4) element stability, transfer or defense (STD); and (5) phage (P) related biological processes (e.g., genome packaging, or lysis and lysogeny). These functions are essential to the persistence of MGEs as independent elements and are orchestrated by an extremely diverse assortment of proteins (16) that we deem suitable as candidate “hallmarks” because of the key roles that they play. Thus, the precise detection of these protein coding genes can serve as the basis for the discovery, classification, and characterization of diverse MGEs in a simple and intuitive way.

## MATERIAL AND METHODS

### Aggregation of a draft pan-mobilome and gene name assignment

A pan-mobilome, i.e., an extensive collection of sequences comprising diverse MGEs, was created by merging sequences produced from seven publicly-available MGE-databases into a single database of protein sequences: ICEBerg 2.0 (13), COMPASS (17), NCBI Plasmid RefSeq Gut Phage Database (18), ISfinder (19), ACLAME (6) and immedb (20). The genomes comprising the basis for the pVOG database (11) and COMPASS, a collection of exclusively nucleotide sequences, were processed with prodigal (v2.6.3) (21) to generate open reading frames using the -p meta setting. The final aggregated dataset included 10,776,849 sequences, or 2,649,813 sequences dereplicated at 97% identity and 80% query coverage (Fig. 1). The 10,776,749 proteins were searched against UniProt (downloaded in the fall of 2020) using diamond blastp, with minimum identity 90% and minimum query coverage of 80% cut-offs. This yielded 8,460,321 matches to UniProt. The 8,460,321 proteins were then used to parse a merged Bacterial, Archaeal, and viral UniProt knowledge base (.dat file) with a custom script (available on the project GitHub page, github.com/clb21565/mobileOG-db/scripts) to extract gene names yielding 110,234 gene names matched to UniProt entries; 20,979 of the 110,234 gene names were unique.

**Figure 1.**
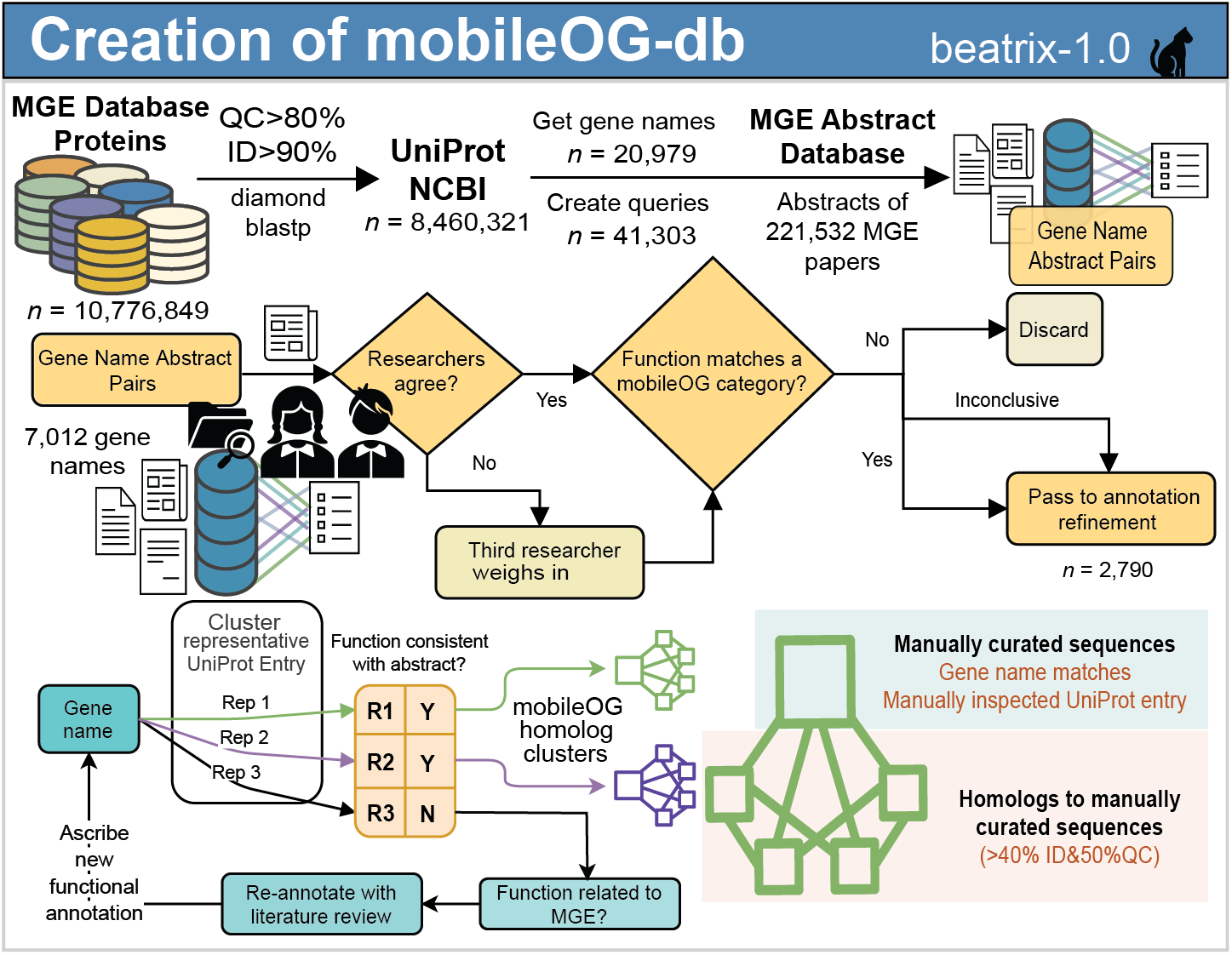
Construction of the mobile orthologous groups database (mobileOG-db). Publicly- available MGE database were downloaded, and their contents mapped to the UniProt TrEMBL/SwissProt knowledge base. Gene names were searched against a filtered database of MGE-related abstracts. 7,012 gene name pairs were then manually inspected by at least two researchers to determine whether the identified gene encoded a protein with a role in one of the target mobileOG categories (replication/recombination/repair, stability/transfer/defence, integration/excision, phage, transfer). A total of 2,790 manually-curated gene names were passed to annotation refinement, where names were paired with UniProt/NCBI entries and associated metadata. To reduce the number of manual curation events needed, we selected one representative sequence for each cluster (40% identity over 50% of reference length using mmseqs2) with a given gene name and compared their database-derived putative function with literature descriptions of the proteins recovered from the abstract analysis. If the UniProt/NCBI entry did not support a link between the gene name and function, the protein was annotated with a literature review.

### Manual curation and annotation of the mobileOG-db

The 20,979 unique gene names were augmented to prepare queries for searching against the MGE abstract database (Supplementary Methods, Table S1). The MGE abstract database was searched using the unique queries and a resulting 8,372 gene name-abstract pairs were manually inspected by at least two researchers. Sequences were manually curated and provided a functional annotation by comparing the abstract text to the putative function reported within each UniProt or NCBI entry (Fig 1). Because the same gene name can be attributed to multiple UniProt entries, sequence entities were annotated on the basis of their 40% identity 50% query/subject coverage (mmseqs easy-clust -c 0) cluster. If the cluster representative had a putative function inconsistent with the attributing abstract(s) (Supplementary Methods), the sequence was reannotated with a review of literature recovered by searching for the gene name and putative function in PubMed. To improve coverage, keyword matches in the fasta headers with a table of MGE protein keywords (Table S2) was used as evidence for inclusion in mobileOG-db. The evidence used to determine inclusion in mobileOG-db (manual curation, homology, or keyword searches) is recorded in mobileOG-db. Examples of our rationale are provided in Supplementary Methods. Last, sequences with matches to SwissProt entries were considered a special case and were manually curated regardless of whether they were returned during the abstract analysis. The gene names, queries, and the abstract database, are available at the FigShare project (https://doi.org/10.6084/m9.figshare.15170736).

## RESULTS

### Catalogue of the mobile orthologous groups database

mobileOG-db consists of five major functional categories (P, RRR, STD, T, and IE) and numerous minor categories, providing intuitive interpretation of search and filter terms. Starting with a pan-mobilome of 10,776,213 proteins derived from ICEBerg, ACLAME, NCBI Plasmid RefSeq, COMPASS, immedb, and ISfinder, we identified proteins performing the defining functions of phages, IGEs, plasmids, and insertion sequences. Owing to the extensive curation effort, a key advance achieved in the present database is its delineation of major and minor mobileOG categories that compose complex elements (Fig. 2A). For example, the *Shigella flexneri* plasmid R100 is displayed with different functional modules coloured by our mobileOG categories (Fig. 2A). There is a prominent RRR module (including *repA*); T: conjugation module (including *finO*); and two transposons (Tn*21* and Tn*10*) dense with IE module protein-coding genes. Altogether, this first release of mobileOG-db (beatrix) comprises 823,797 dereplicated proteins including over 29,000 derived directly from manually curated entries; 6,140 protein clusters or families (defined as greater than 40% identical over 50% of the subject and query length, see methods); 2,444 unique manual annotations, and 1,393 references (Fig. 2B).

**Figure 2.**
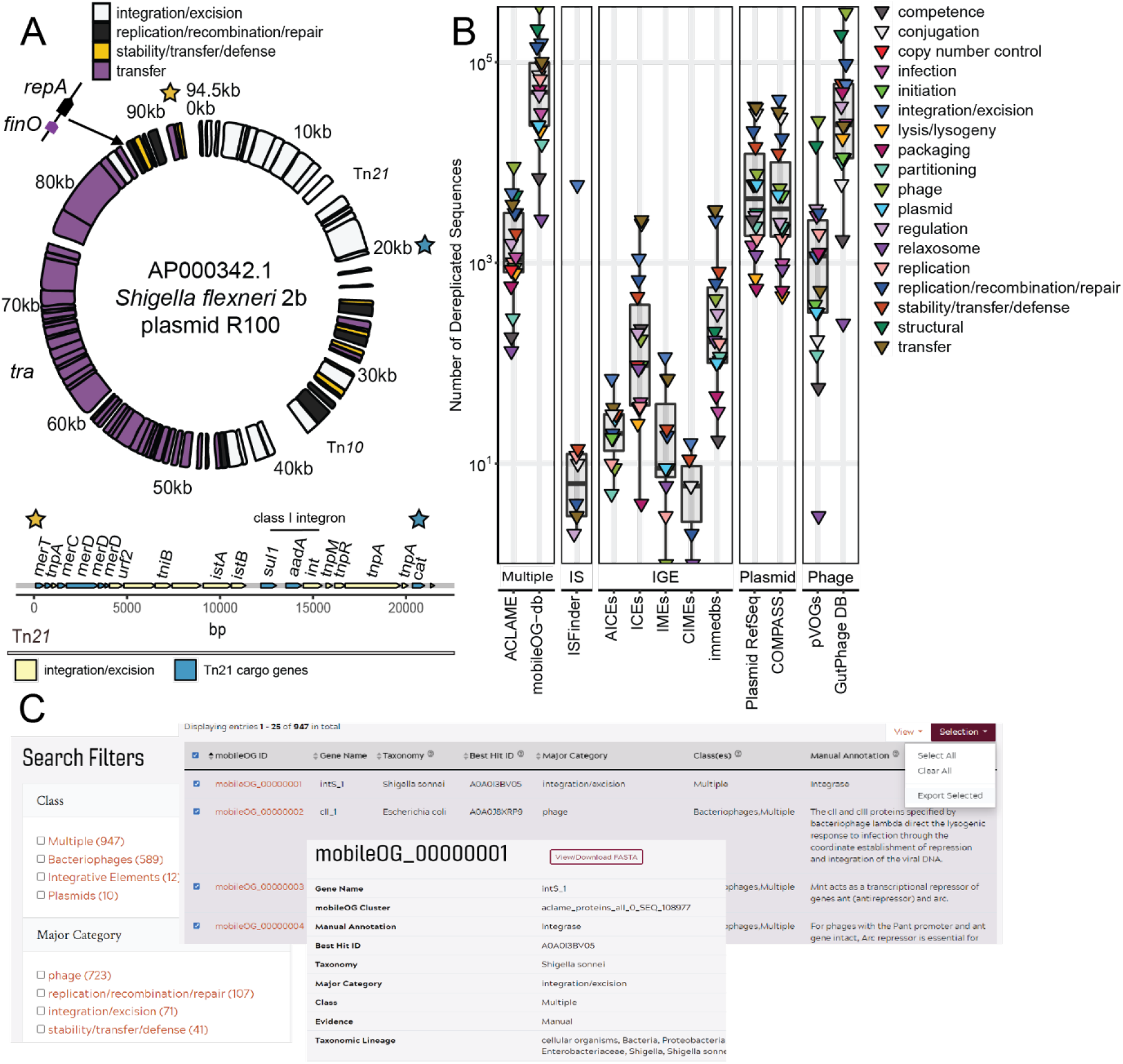
Overview and examples of the content within mobileOG-db. (A) Shigella flexneri plasmid R100, among the first conjugative multidrug resistance plasmids to be identified carries Tn21, a resistance geneharbouring transposon (initial and terminal repeats indicated by gold and blue stars, respectively). Usage of the mobileOG-db positively identifies AP000342.1 as a conjugative plasmid enriched with integration/excision module proteins highlighting the carriage of integrative elements IS1353, IS1326, IS1b, and a class 1 integron. In the bottom panel, Tn21 is examined in greater detail; Tn21 cargo genes (blue) are not included in mobileOG-db. (B) mobileOG-db contains a substantially improved diversity of sequences compared to presently available databases, covering a wide-array of MGE functional modules. Abbreviations: IS: Insertion sequences recovered from ISfinder.; IGE: integrative elements recovered from ICEberg. (C) the mobileOG-db webserver provides a user-friendly platform for interactive annotation database customization for scientists across the life sciences. Researchers can search, browse, filter, and download customized datasets tailored to their research questions.

### Usage recommendations and examples

For detecting and classifying elements from long genomic segments (i.e., long reads or assembled short reads), it is recommended that an accurate annotation consists of multiple co-localized hits in close proximity, similar to the pattern-based co-localization approach leveraged by ICEBerg (5) for IGE discovery. Likewise, it is noted that hits solely to RRR modules are not necessarily indicative of an MGE; plasmids and phages frequently encode homologs of RRR machinery (22) that are also present in exclusively cellular DNA. An additional caveat is that hits to type four secretion systems may not be indicative of a MGE; paralogues of these proteins are also virulence determinants in some organisms (23). A preliminary annotation pipeline has been developed (Supplementary Methods, Table S1) to allow for automated element annotation (Supplementary Methods). Usage of mobileOG-db enabled successful classification of up to 98.2% and 99.7% of the plasmids and phages, respectively, comprising a test dataset (https://doi.org/10.6084/m9.figshare.15170736. Table S3) of genomes in the COMPASS or the GutPhage databases. Other uses might include the creation of quantitative metrics for horizontal gene transfer hypothesis testing (Fig S3).

The mobileOG-db web portal provides a user-friendly interface for scientists across relevant fields intersecting the Life Sciences to browsing and customizing datasets of MGE proteins (Fig. 2C). Usage of the website allows users to select different major and minor mobileOG categories to hone their experimental design or intended usage. Further, the ability to filter and select from different element-types allow for the user to identify genes that occur across different element types. For instance, users could select experimentally validated insertion sequence proteins that also occur on plasmids, a key route for the horizontal transmission of ARGs (24).

## DISCUSSION

The creation of mobileOG-db was motivated by a lack of an up-to-date and comprehensive resource for markers of diverse classes of MGEs. Here, using a layered annotation scheme, we analysed 10,776,212 MGE-encoded proteins to differentiate sequences that are anticipated to be informative for annotation from those that are not defensible for this purpose from a biological standpoint. Importantly, we recognize that the annotation framework implemented here cannot produce a highly granular description of MGE function. However, providing such a resource for every major class of bacterial MGEs is far beyond the scope of the present work and, in addition to uncertainty of protein function, element-specific resources are available that form the basis for mobileOG-db. Instead, mobileOG-db provides a “Swiss Army knife” that can serve as the foundation for an array of analyses, which can be designed, customized, and refined using the web service.

Looking towards the future, the delineation of MGE functional modules and conserved protein families could potentially support probabilistic methods for clustering, annotating, and classifying MGEs. Such frameworks could also provide a basis for other analyses leveraging compositional and structural features of the elements to quantitatively estimate potential host-linkages, cargo, and the potential for transmission to clinically relevant pathogens. These analyses are being developed for inclusion in a future release of mobileOG-db and show promise for harnessing large scale genomic data for predictive public health insights.

## DATA AVAILABILITY

mobileOG-db is available at mobileogdb.flsi.cloud.vt.edu/, where users can browse, filter, search, and download customized datasets and references. Scripts used in the text mining analysis and two example pipelines using R or Python are available at https://github.com/clb21565/mobileOG-db/scripts.

## SUPPLEMENTARY DATA

Supplementary Data are available at NAR online and the manuscript FigShare repository https://doi.org/10.6084/m9.figshare.15170736.

## ACKNOWLEDGEMENT

We would like to directly express our appreciation for the work and expertise that went into designing the databases making up mobileOG-db. The authors acknowledge the Advanced Research Computing at Virginia Tech for providing computational resources.

## REFERENCES

1. Rankin, D.J., Rocha, E.P.C. and Brown, S.P. (2011) What traits are carried on mobile genetic elements, and why. Heredity (Edinb)., 106, 1–10.

2. Berg, O.G. and Kurland, C.G. (2002) Evolution of microbial genomes: Sequence acquisition and loss. Mol. Biol. Evol., 19, 2265–2276.

3. Partridge, S.R., Kwong, S.M., Firth, N. and Jensen, S.O. (2018) Mobile genetic elements associated with antimicrobial resistance. Clin. Microbiol. Rev., 31.

4. Ellabaan, M.M.H., Munck, C., Porse, A., Imamovic, L. and Sommer, M.O.A. (2021) Forecasting the dissemination of antibiotic resistance genes across bacterial genomes. Nat. Commun., 12, 1–10.

5. Liu, M., Li, X., Xie, Y., Bi, D., Sun, J., Li, J., Tai, C., Deng, Z. and Ou, H.Y. (2019) ICEberg 2.0: An updated database of bacterial integrative and conjugative elements. Nucleic Acids Res., 47, D660–D665.

6. Leplae, R., Lima-Mendez, G. and Toussaint, A. (2009) ACLAME: A CLAssification of mobile genetic elements, update 2010. Nucleic Acids Res., 38.

7. Roux, S., Enault, F., Hurwitz, B.L. and Sullivan, M.B. (2015) VirSorter: Mining viral signal from microbial genomic data. PeerJ, 2015, e985.

8. Krawczyk, P.S., Lipinski, L. and Dziembowski, A. (2018) PlasFlow: predicting plasmid sequences in metagenomic data using genome signatures. Nucleic Acids Res., 46, e35.

9. Orlek, A., Stoesser, N., Anjum, M.F., Doumith, M., Ellington, M.J., Peto, T., Crook, D., Woodford, N., Sarah Walker, A., Phan, H., et al. (2017) Plasmid classification in an era of whole-genome sequencing: Application in studies of antibiotic resistance epidemiology. Front. Microbiol., 8, 1–10.

10. Carr, V.R., Shkoporov, A., Hill, C., Mullany, P. and Moyes, D.L. (2021) Probing the Mobilome: Discoveries in the Dynamic Microbiome. Trends Microbiol., 29, 158–170.

11. Grazziotin, A.L., Koonin, E. V. and Kristensen, D.M. (2017) Prokaryotic Virus Orthologous Groups (pVOGs): A resource for comparative genomics and protein family annotation. Nucleic Acids Res., 45, D491–D498.

12. Pfeifer, E., Moura De Sousa, J.A., Touchon, M. and Rocha, E.P.C. (2021) Bacteria have numerous distinctive groups of phage-plasmids with conserved phage and variable plasmid gene repertoires. Nucleic Acids Res., 49, 2655–2673.

13. Liu, M., Li, X., Xie, Y., Bi, D., Sun, J., Li, J., Tai, C., Deng, Z. and Ou, H.Y. (2019) ICEberg 2.0: An updated database of bacterial integrative and conjugative elements. Nucleic Acids Res., 47, D660–D665.

14. Slizovskiy, I.B., Mukherjee, K., Dean, C.J., Boucher, C. and Noyes, N.R. (2020) Mobilization of Antibiotic Resistance: Are Current Approaches for Colocalizing Resistomes and Mobilomes Useful? Front. Microbiol., 11, 1376.

15. Partridge, S.R., Kwong, S.M., Firth, N. and Jensen, S.O. (2018) Mobile genetic elements associated with antimicrobial resistance. Clin. Microbiol. Rev., 31.

16. Craig, N.L. (2015) A Moveable feast: An introduction to mobile DNA. In Mobile DNA III. wiley, pp. 3–39.

17. Douarre, P.E., Mallet, L., Radomski, N., Felten, A. and Mistou, M.Y. (2020) Analysis of COMPASS, a New Comprehensive Plasmid Database Revealed Prevalence of Multireplicon and Extensive Diversity of IncF Plasmids. Front. Microbiol., 11, 483.

18. Camarillo-Guerrero, L.F., Almeida, A., Rangel-Pineros, G., Finn, R.D. and Lawley, T.D. (2021) Massive expansion of human gut bacteriophage diversity. Cell, 184, 1098-1109.e9.

19. Siguier, P., Perochon, J., Lestrade, L., Mahillon, J. and Chandler, M. (2006) ISfinder: the reference centre for bacterial insertion sequences. Nucleic Acids Res., 34.

20. Jiang, X., Hall, A.B., Xavier, R.J. and Alm, E.J. (2019) Comprehensive analysis of chromosomal mobile genetic elements in the gut microbiome reveals phylum-level niche-adaptive gene pools. PLoS One, 14, e0223680.

21. Hyatt, D., Chen, G.L., LoCascio, P.F., Land, M.L., Larimer, F.W. and Hauser, L.J. (2010) Prodigal: Prokaryotic gene recognition and translation initiation site identification. BMC Bioinformatics, 11, 119.

22. Chen, S.H., Byrne, R.T., Wood, E.A. and Cox, M.M. (2015) Escherichia coli radD(yejH) gene: A novel function involved in radiation resistance and double-strand break repair. Mol. Microbiol., 95, 754–768.

23. Costa, T.R.D., Harb, L., Khara, P., Zeng, L., Hu, B. and Christie, P.J. (2021) Type IV secretion systems: Advances in structure, function, and activation. Mol. Microbiol., 115, 436–452.

24. Che, Y., Yang, Y., Xu, X., Brinda, K., Polz, M.F., Hanage, W.P. and Zhang, T. (2021) Conjugative plasmids interact with insertion sequences to shape the horizontal transfer of antimicrobial resistance genes. Proc. Natl. Acad. Sci. U. S. A., 118.

