## Supplemental Information and Data for "mobileOG-db: a manually curated database of protein families mediating the life cycle of bacterial mobile genetic elements"

### SUPPLEMENTAL METHODS

(i) Example rationale for annotating proteins
(ii) Description of the mobileOG-kyanite for autonomous element detection and classification

### SUPPLEMENTARY DATA

**Table S1.** Keywords used to identify mobile genetic element abstracts in PubMed.

**Table S2.** Table of keywords and their associated categories created to identify putative MGE sequences in the merged database that are associated with the target categories.

**Table S3.** Evaluation of mobileOG-kyanite, a pipeline for identifying putative mobile element contigs. Attached as csv.

**Figure S1.** Description of mobileOG.pl-kyanite, a preliminary pipeline for autonomous element detection and classification.

**Figure S2.** Comparative analysis of GC content of element “core” genes allows for horizontal gene transfer hypothesis testing**.** Color scheme is fold change of GC content of integrative element (IGE) genes relative to the “core” mobileOG-db annotated genes.

### Supplemental Methods

(I) Creation and querying of a mobile genetic element abstract database.

A list of descriptive keywords related to MGEs (Table S1) was developed to identify a comprehensive selection of MGE-related literature for use in text-mining aided manual curation of MGE-protein function. These terms were searched against NCBI PubMed using entrez to extract article metadata and abstracts. To weed out false-positive hits, the preliminary set of abstracts were then refined to exclude those with less than 3 keyword matches. The final resulting abstract were then used in subsequent searches and manual curation.

Queries were constructed in the following way. Gene names with three or fewer characters were prefixed with “protein” or “gene” to reduce spurious hits. Gene names with an underscore or special character were split on either side (e.g., *tnpA_2* would become *tnpA* and *2*) and both sides were used as queries against the MGE-abstract database. Altogether this produced 41,303 unique queries (the complete list of search terms is available on the figshare project site: <https://doi.org/10.6084/m9.figshare.15170736>).

(II) Example rationale of protein annotations.

Proteins were included in mobileOG-db only if there was experimental evidence of their direct involvement with one of the targeted functions. Proteins with only indirect interactions with one of the target functions were not included unless they had been shown to be essential for element persistence or replication. For example, these criteria excluded ribonucleotide reductases found within many phage genomes (1), which only have an indirect impact on replication through nucleotide metabolism (2, 3), except under conditions of anaerobic growth (2, 4). While these proteins are useful indicators of phage diversity (5, 6), we were unable to find evidence of a direct role in replication other than nucleotide metabolism and thus these proteins are not present in mobileOG-db. By contrast, phage-encoded thymidylate synthase homologs provide nucleotide substrates for replication and control levels of methyl- or hydroxymethyl- thymidine monophosphates (7). These modified pyrimidines can then be further hypermodified (8) by additional functional moieties (7, 9), which alter the steric properties of the nucleic acid of the viral genome. This process can therefore provide a phage genome with defense against host-encoded CRISPR (10) and restriction modification systems (11–14). Thus, thymidylate synthases were included in mobileOG-db and categorized in the replication/recombination/repair major category with minor categories stability and defense.

By contrast, we found that there were several examples of proteins with names that did not match the results of the abstract database, and therefore had to be manually curated to reconcile the disagreement. For example,

tr|A0A2Z2Q3C7|A0A2Z2Q3C7_9RHIZ Polyamine ABC transporter ATP-binding protein OS=Agrobacterium larrymoorei OX=160699 GN=repB PE=3 SV=1

The protein repB was identified as a regulator of plasmid replication by the abstract analysis and this sequence initially appeared to be an erroneous attribution of the name, or a protein with the same name but different function. Upon further inspection, it became apparent that the header was not descriptive of the putative function of the protein:

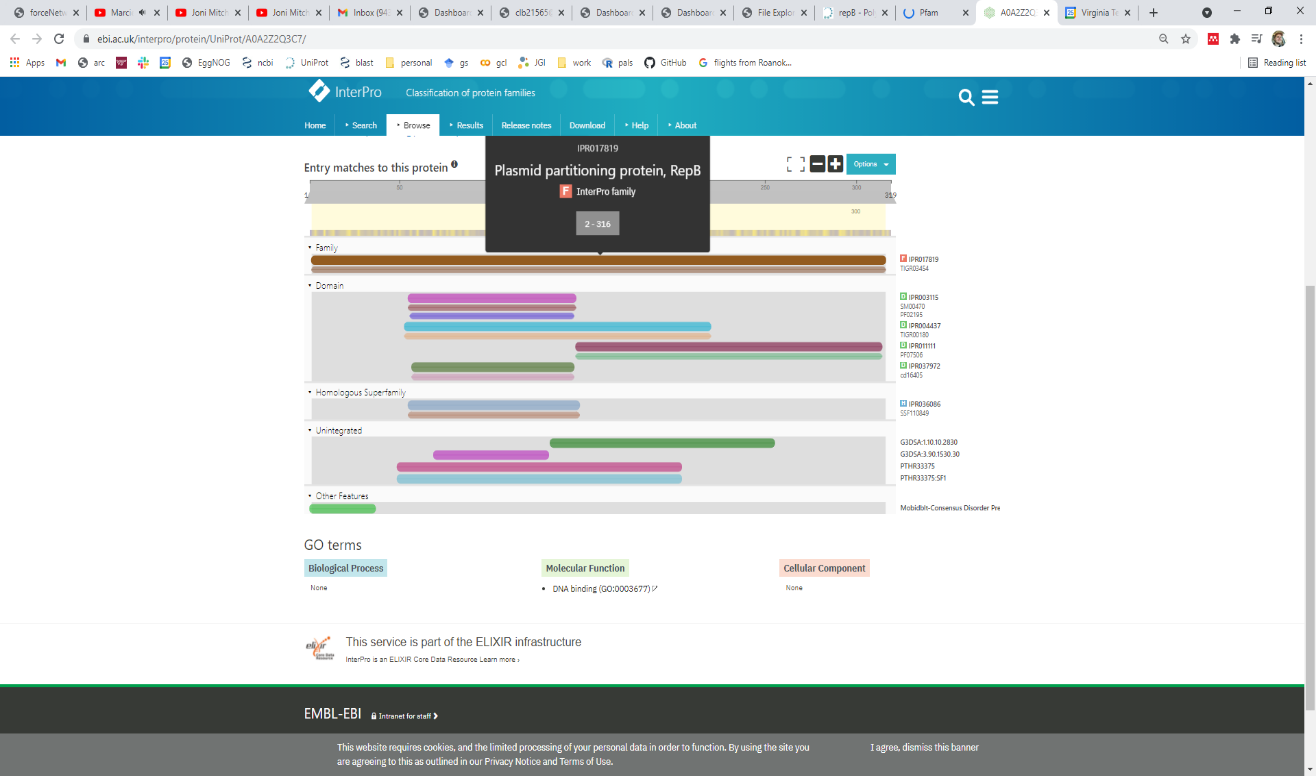
 [View protein in InterPro](https://www.ebi.ac.uk/interpro/protein/A0A2Z2Q3C7)
[IPR004437](https://www.ebi.ac.uk/interpro/entry/IPR004437), ParB/RepB/Spo0J
[IPR003115](https://www.ebi.ac.uk/interpro/entry/IPR003115), ParB/Sulfiredoxin_dom
[IPR036086](https://www.ebi.ac.uk/interpro/entry/IPR036086), ParB/Sulfiredoxin_sf
[IPR017819](https://www.ebi.ac.uk/interpro/entry/IPR017819), Plasmid_partition_RepB
[IPR011111](https://www.ebi.ac.uk/interpro/entry/IPR011111), Plasmid_RepB
[IPR037972](https://www.ebi.ac.uk/interpro/entry/IPR037972), RepB_N

**Figure S1.** Example of incorrect annotation manually reconciled in mobileOG-db.

Thus, this entry was included in the manually curated sequences as it had a positive association between name, literature, and putative function. UniProt was additionally contacted to seek a correction for this entry.

Below are two examples of MGE gene names that also correspond to names of other genes and proteins. *mobC* is also the name of a gene encoding a mobilase associated with conjugal plasmid transfer (15); *motA* also refers to a gene encoding a T4 phage transcriptional regulator (16).

tr|A0A0K2CS33|A0A0K2CS33_CITFR Molybdopterin-guanine dinucleotide biosynthesis protein mobc OS=Citrobacter freundii OX=546 GN=mobC PE=4 SV=1

tr|A0A174YTE7|A0A174YTE7_9FIRM Chemotaxis protein MotA OS=[Eubacterium] eligens OX=39485 GN=motA PE=4 SV=1

(ii)	mobileOG-db.pl-kyanite, a preliminary pipeline to detect and classify genomic contigs or long reads as putative MGEs.

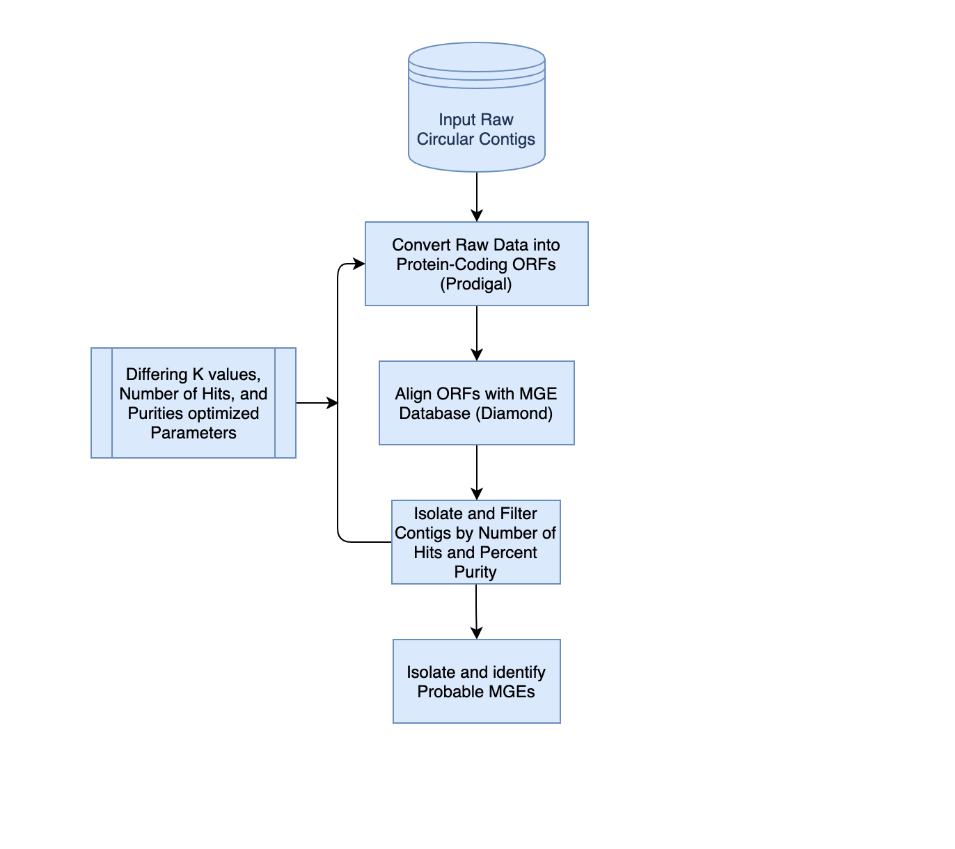

**Figure S2.** mobileOG-db.pl-kyanite takes genomic contigs as input, converts the nucleotide sequences to open reading frames using prodigal, then aligns the open reading frames against mobileOG-db. Different diamond settings can be used, and were tested for recovering phages or plasmids from a test data set.

### SUPPLEMENTARY DATA

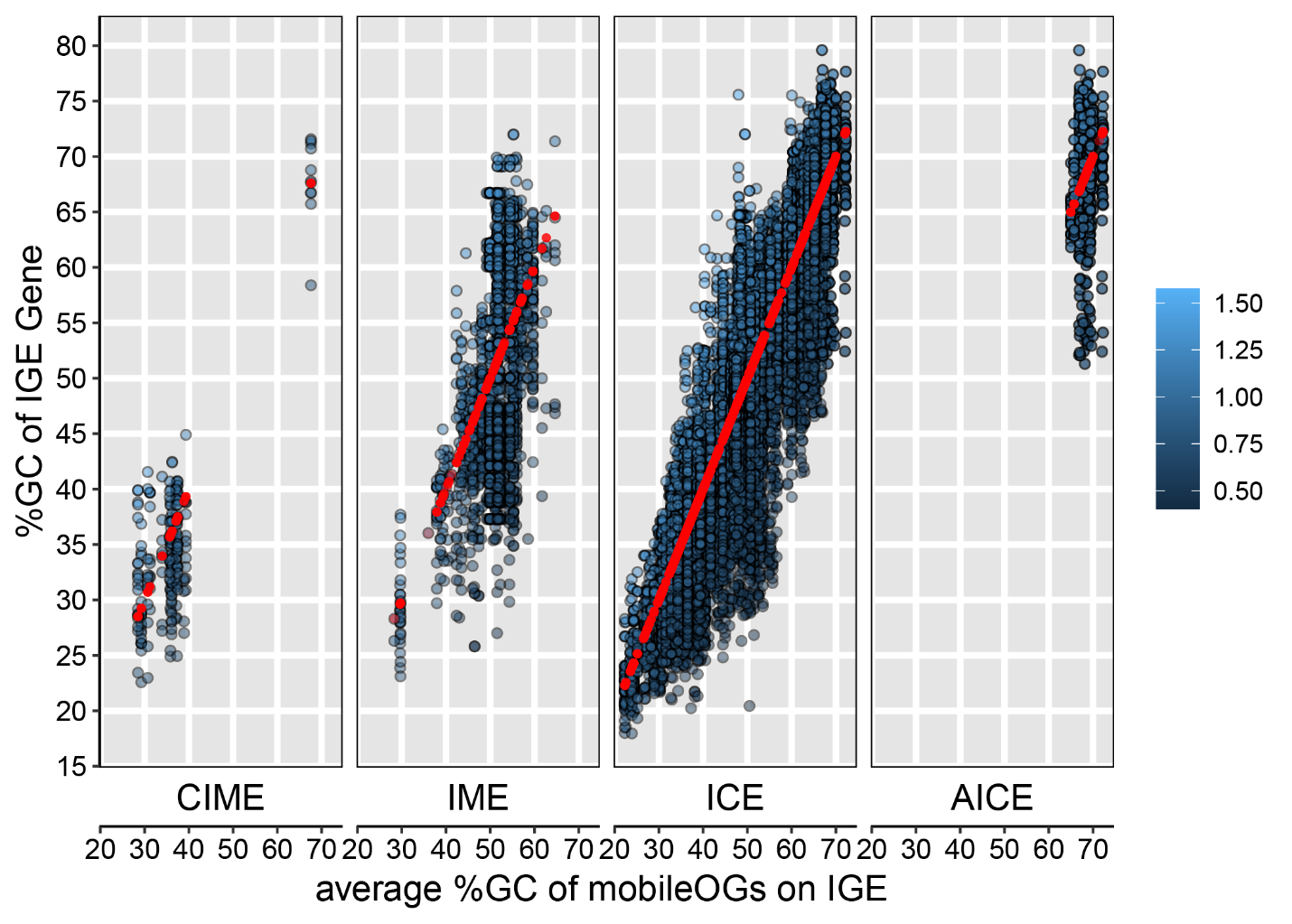

**Figure S3.** Comparative GC% analysis allows for the computation of a biologically reasoned quantitative metric for test hypothesis around horizontal gene transfer. Blue scale is fold-difference between the GC% of a gene relative to the average GC% of the mobileOGs present on the element carrying it.

### SUPPLEMENTARY DATA

Table S1. Keywords used to identify MGE abstracts.

Keyword

competence

CRISPR

nuclease

replication

toxin

antitoxin

addiction

transposition

replication

DNA

capsid

tape measure

terminase

tail collar

baseplate

Reverse transcriptase

resolvase

invertase

shufflon

restriction

methyltransferase

mobile genetic element

transposon

integrative conjugative element

chromosomal integrative mobile element

mobile DNA

virus

prophage

phage

plasmid

incompatibility group

mobile

selfish genetic element

casposon

viral

proviral

insertion sequence

restriction modification

pINC

ICEBerg

mobilome

excision

integration

recombination

transposable element

| Table S2. Keywords used to recover MGE functional proteins from the merged database. | | |
| --- | --- | --- |
| Category | Include | Do not include |
| phage,structural | head,neck,capsid,baseplate,vertex,whisker,tail, sheathe,portal,coat,spike,neck,tape measure,virion,base plate,Tape-measure,Plate protein | conjugation,type VI secretion system,cytochrome c oxidase,two-tailed,cluster,conjugal,photosystem II stability,hammerhead,pilus,conjugative |
| phage,lysogeny | lysin,autolysin,endolysin,lysozyme,holin,antiholin,spanin,abortive infection | lysozyme if no "phage" or "virus"; hemolysin, haemolysin,choline,Lysinibacillus, hydrolysing |
| phage,regulation | regulatory cii,prophage repressor,tapemeasure,antirepressor,anti-repressor,phage late control |  |
| phage,replication,packaging | terminase,terl | interleukin |
| integration,excision | integration,excision,integrase,tyrosine recombinase,serine recombinase,serine integrase,phage integrase,transposase,helper of transposition,excisionase,xis protein,cassette chromosome recombinase,Integration host factor,recombination directionality factor,shufflon,group I intron endonuclease,Tnp domain,Retron-type reverse transcriptase,intron endonuclease | chemotaxis |
| integration,excision,inversion | invertase,inversion |  |
| integration,excision,replication,recombination,repair | resolvase |  |
| stability,transfer,defense | addiction,toxin/antitoxin,antitoxin,YoeB,YoeF,HigB,CRISPR,toxin-antitoxin,RelE/ParE,entry exclusion,stbB,plasmid stabilization system,DNA methylase,restriction endonuclease,surface exclusion,restriction-modification,Protein kilB,kilB,Hok/Gef ,N-6-adenine-methyltransferase,N-6 DNA methylase,restriction enzyme,DNA adenine methylase | shiga toxin,Clavibacter michiganensis,michiganensis,RIGHA |
| transfer,conjugation | conjugation,pilus,conjugational,conjugative,type IV secretion system protein,mobilization,relaxase,mobilase,FtsK/SpoIIIE,FtsK,SpoIIIE,TraB,TraM,conjugal,VirB3,MobA/MobL,TrbC/VirB2 | tram |
| CRISPR | CRISPR |  |
| transfer,competence | competence |  |
| replication,regulation | protein RepA, repZ, repL, |  |
| phage,infection | adsorption,antireceptor,Super-infection exclusion |  |
| replication,transfer,partitioning | ParB,RepB,Spo0J |  |
| transfer | DNA transfer |  |

The “include” column records which terms were searched for, and the do not include column records the search terms used to filter (remove) erroneous hits to a given category following the search.

Supplementary References

1. Iyer,L.M., Anantharaman,V., Krishnan,A., Burroughs,A.M. and Aravind,L. (2021) Jumbo Phages: A Comparative Genomic Overview of Core Functions and Adaptions for Biological Conflicts. *Viruses*, **13**.

2. Lundin,D., Torrents,E., Poole,A.M. and Sjöberg,B.-M. (2009) RNRdb, a curated database of the universal enzyme family ribonucleotide reductase, reveals a high level of misannotation in sequences deposited to Genbank. *BMC Genomics 2009 101*, **10**, 1–8.

3. ES,M., E,K., G,M., F,A., T,K. and W,R. (2003) Bacteriophage T4 genome. *Microbiol. Mol. Biol. Rev.*, **67**, 86–156.

4. Fontecave,M., Mulliez,E. and Logan,D.T. (2002) Deoxyribonucleotide synthesis in anaerobic microorganisms: The class III ribonucleotide reductase. *Prog. Nucleic Acid Res. Mol. Biol.*, **72**, 95–127.

5. B,D., B,X., D,L., RA,E. and M,B. (2013) A bioinformatic analysis of ribonucleotide reductase genes in phage genomes and metagenomes. *BMC Evol. Biol.*, **13**.

6. Sakowski,E.G., Munsell,E. V., Hyatt,M., Kress,W., Williamson,S.J., Nasko,D.J., Polson,S.W. and Wommack,K.E. (2014) Ribonucleotide reductases reveal novel viral diversity and predict biological and ecological features of unknown marine viruses. *Proc. Natl. Acad. Sci. U. S. A.*, **111**, 15786–15791.

7. Weigele,P. and Raleigh,E.A. (2016) Biosynthesis and Function of Modified Bases in Bacteria and Their Viruses. 10.1021/acs.chemrev.6b00114.

8. JH,G.-A. and P,B. (1995) Hypermodified bases in DNA. *FASEB J.*, **9**, 1034–1042.

9. YJ,L., N,D., SE,W., S,M., ME,F., KM,K., C,G., IR,C. and PR,W. (2018) Identification and biosynthesis of thymidine hypermodifications in the genomic DNA of widespread bacterial viruses. *Proc. Natl. Acad. Sci. U. S. A.*, **115**, E3116–E3125.

10. Bryson,A.L., Hwang,Y., Sherrill-Mix,S., Wu,G.D., Lewis,J.D., Black,L., Clark,T.A. and Bushman,F.D. (2015) Covalent modification of bacteriophage T4 DNA inhibits CRISPRCas9. *MBio*, **6**.

11. Flodman,K., Tsai,R., Xu,M.Y., Corrêa,I.R., Jr., Copelas,A., Lee,Y.-J., Xu,M.-Q., Weigele,P. and Xu,S. (2019) Type II Restriction of Bacteriophage DNA With 5hmdU-Derived Base Modifications. *Front. Microbiol.*, **10**, 584.

12. PB,M., WW,W. and RA,W. (1985) alpha-Putrescinylthymine and the sensitivity of bacteriophage phi W-14 DNA to restriction endonucleases. *Nucleic Acids Res.*, **13**, 2559–2568.

13. Huang,L.H., Farnet,C.M., Ehrlich,K.C. and Ehrlich,M. (1982) Digestion of highly modified bacteriophage DNA by restriction endonucleases. *Nucleic Acids Res.*, **10**, 1579.

14. DH,K. and TA,B. (1983) Bacteriophage survival: multiple mechanisms for avoiding the deoxyribonucleic acid restriction systems of their hosts. *Microbiol. Rev.*, **47**, 345–360.

15. Garcillán-Barcia,M.P., Francia,M.V. and De La Cruz,F. (2009) The diversity of conjugative relaxases and its application in plasmid classification. In *FEMS Microbiology Reviews*.Vol. 33, pp. 657–687.

16. Schmidt,R.P. and Kreuzer,K.N. (1992) Purified MotA protein binds the -30 region of a bacteriophage T4 middle-mode promoter and activates transcription in vitro. *J. Biol. Chem.*, **267**, 11399–11407.
